# Aridity modulates biogeographic distribution and community assembly of cyanobacterial morphotypes in drylands

**DOI:** 10.1101/2022.05.30.494084

**Authors:** Xiaoyu Guo, Hua Li, Da Huo, Renhui Li, Silong Zhang, Chunxiang Hu, Gongliang Yu, Lirong Song

## Abstract

The patterns of biogeographical distribution and assembly processes of microbiota are of vital importance for understanding ecological adaptation and functioning maintenance that microorganisms provide. Although the assembly of microbial communities was increasingly inspected, the role of their morphological characteristics is still poorly ascertained. Here, by integrating high-throughput sequencing and traditional trait-based classification, we investigated taxonomic and phylogenetic turnovers of different morphotypes of terrestrial cyanobacteria in biocrusts, as a model system, to evaluate the contributions of deterministic and stochastic processes across a large scale in drylands. The results showed that the bundle-forming category dominated in arid ecosystems with high tolerance against environmental fluctuations. Despite strong distance-decay relationships of *β*-diversity in all categories, both spatial composition and phylogenetic turnover rates of bundle-forming cyanobacteria were significantly lower than unicellular/colonial, heterocystous, and other non-heterocyst filamentous cyanobacteria. Null model analysis of phylogenetic signals and abundance-based neutral model found that stochastic processes prevailed in the assembly, while aridity mediated the balance between determinism and stochasticity and prompted a shifting threshold among morphotypes. Our findings provide a unique perspective to understand the role of microbial morphology, highlighting the differentiation of biogeographic distribution, environmental response, and species preference between morphotypes with consideration of potential functional consequences, and therefore facilitated the prediction of biodiversity loss under climate change.

## 1. Introduction

Microorganisms in soils constitute an immensely diverse consortium on Earth [1, 2]. They represent one of the most complex communities that play crucial roles in biogeochemical cycling and contribute greatly to multiple functions of terrestrial ecosystems [3-5]. Evidence is mounting that community characteristics, such as richness, composition, interaction and dynamics are particularly essential for microbial functionality [4, 6, 7]. Thus, unraveling the underlying mechanisms driving community assembly across spatiotemporal scales has become increasingly a central but still challenging task for microbial ecologists to understand the functional consequences of biodiversity loss in soil microbiota [8, 9].

Although the factors controlling the patterns of microbial diversity remain highly controversial, a growing consensus is that deterministic, as well as stochastic, processes collectively regulate the assembly of local communities and biogeographic distribution [10, 11]. The set of deterministic factors, derived from the traditional niche-based hypothesis, includes both abiotic conditions (*e*.*g*., habitat heterogeneity or environmental gradients) and biotic variables (*e*.*g*., species traits and interactions) [11, 12]. However, determinism alone is not sufficient to depict the entire variation of microbial communities [13, 14]. Numerous studies have demonstrated that the stochastic processes of random disturbance, dispersal, extinction, and speciation could be equally, if not more, important in governing community structures in soils [15-17]. Recently, the balance of determinism versus stochasticity on the microbial assembly was found to be mediated by environmental factors, and the relative contribution could shift largely over space and time [18, 19]. Given the distinct responses of microorganisms [2, 20], whether various subcommunities, categorized by phylogeny, function, or phenotype, are controlled by differential manners of deterministic and stochastic processes, and the extent to which their relative importance explains the variation of community structure, succession, and biogeography needs to be further assessed.

Inspired by Vellend’s conceptual framework [21], a few studies have attempted to quantitatively delineate assembly mechanisms between contrary pairs of subcommunities [22, 23]. For instance, in desert hypolithic communities, the photosynthetic assemblage was subject to stochastic processes, while the heterotrophic assemblage displayed more signatures of niche selection (*i*.*e*., deterministic process) [17]. As yet, albeit emerging studies have elucidated the differences of environmental sensitivity, distribution patterns, and drivers of community assembly in the microbiota [3, 24], the properties of microbial morphology are often neglected, which are supposed to have significant functional signals and modulate the succession of persisting species [25, 26]. A previous study suggested that various bacterial taxa from a particular phylotype to a large extent share similar phenotypic traits or life-history strategies whereby they could adapt to a specific environment [1]. However, it is noteworthy that a majority of testifies on the roles of morphological traits prefer to focus on vascular plant communities [27-29], heavily due to the lack of conservatively observable features of microbial morphotypes. It restricts efforts to disentangle the biogeographic patterns and mechanisms of community assembly to predict the response of soil microorganisms to climate change [30].

Nevertheless, terrestrial cyanobacteria in biological soil crusts (biocrusts), as pioneers on nutrient-poor and abiotically stressful substrates [31], are a unique minority of microbes that exert high differentiation of morphological characteristics to create stable organic-rich layers on topsoil [32, 33]. In global drylands, biocrusts constitute up to 70% of the soil surface [34], providing critical ecosystem functions, such as water retention, nutrient accumulation, and soil stabilization [20]. Many studies have underlined the role of terrestrial cyanobacteria as key biological agents in the remediation and promotion of soil environments [35, 36].

According to the well-differentiated morphological properties, terrestrial cyanobacteria are proposed to be categorized into four main morphotypes, including bundle-forming cyanobacteria that dominate arid/semiarid environments worldwide (*e*.*g*., *Microcoleus* spp.), other non-heterocyst filamentous cyanobacteria (*e*.*g*., *Leptolyngbya* spp.), heterocystous cyanobacteria belonging to Nostocales that can fix atmospheric nitrogen (*e*.*g*., *Scytonema* spp.), and unicellular/colonial cyanobacteria (*e*.*g*., *Chroococcidiopsis* spp.) [37, 38]. It has been demonstrated that cyanobacterial functionalities in biocrusts, as well as their adaptation to abiotic stresses, rely on the performance of various morphological characteristics [39-41]. The response difference of cyanobacteria against ambient fluctuation may impact compositional turnover and biogeographic distribution, generating potential functional alterations. However, little is known about whether and how morphotypes of terrestrial cyanobacteria in biocrusts assemble in distinct manners consistent with environmental heterogeneity.

To address these concerns, we quantitatively detected the biogeographic features and processes of community assembly in four cyanobacterial morphotypes along a natural gradient of northwestern China. The composition of terrestrial cyanobacteria was determined by the high-throughput sequencing method to facilitate empirical identification of morphology. A null model-based evaluation of community assembly was used by calculating phylogenetic and taxonomic turnovers, coupling with an abundance-based neutral model analysis. We then related the environmental variables to species abundance, *β*-diversity, and phylogenetic distribution to reflect ecological preferences of cyanobacterial morphotypes. The results provide a novel perspective on the associations between phenotypic characteristics of microorganisms and biogeographic patterns with a substantial gradient of the environment, and could promote our understanding of the underlying mechanisms of community assembly in soil microbiota.

## 2. Methods

### 2.1 Study sites and sample collection

We collected biocrust samples from 45 sites across the climatic transition belt of northern China in August 2020, spanning a large scale of approximately 37∼41 °N and 100∼110 °E (**Figure 1a, Table S1**). The spatial interval between study sites was at least 1 km apart. The mean annual temperature and precipitation of the sampling regions were 5.4∼9.2 °C and 131∼315 mm, respectively. The topsoil of most sites is covered by biocrusts with sparse high plants. In the open field out of herb/shrub patches, at least three plots (1 m×1 m) were selected randomly to collect the upper biocrust cores (0∼2 cm deep) in triplicate and then homogenized thoroughly. All samples were dried at 37 °C to constant weight in a relatively short period. The air-dried samples were ground and sieved to remove gravel or plant roots, and then stored at −20° for subsequent analyses.

**Fig 1.**
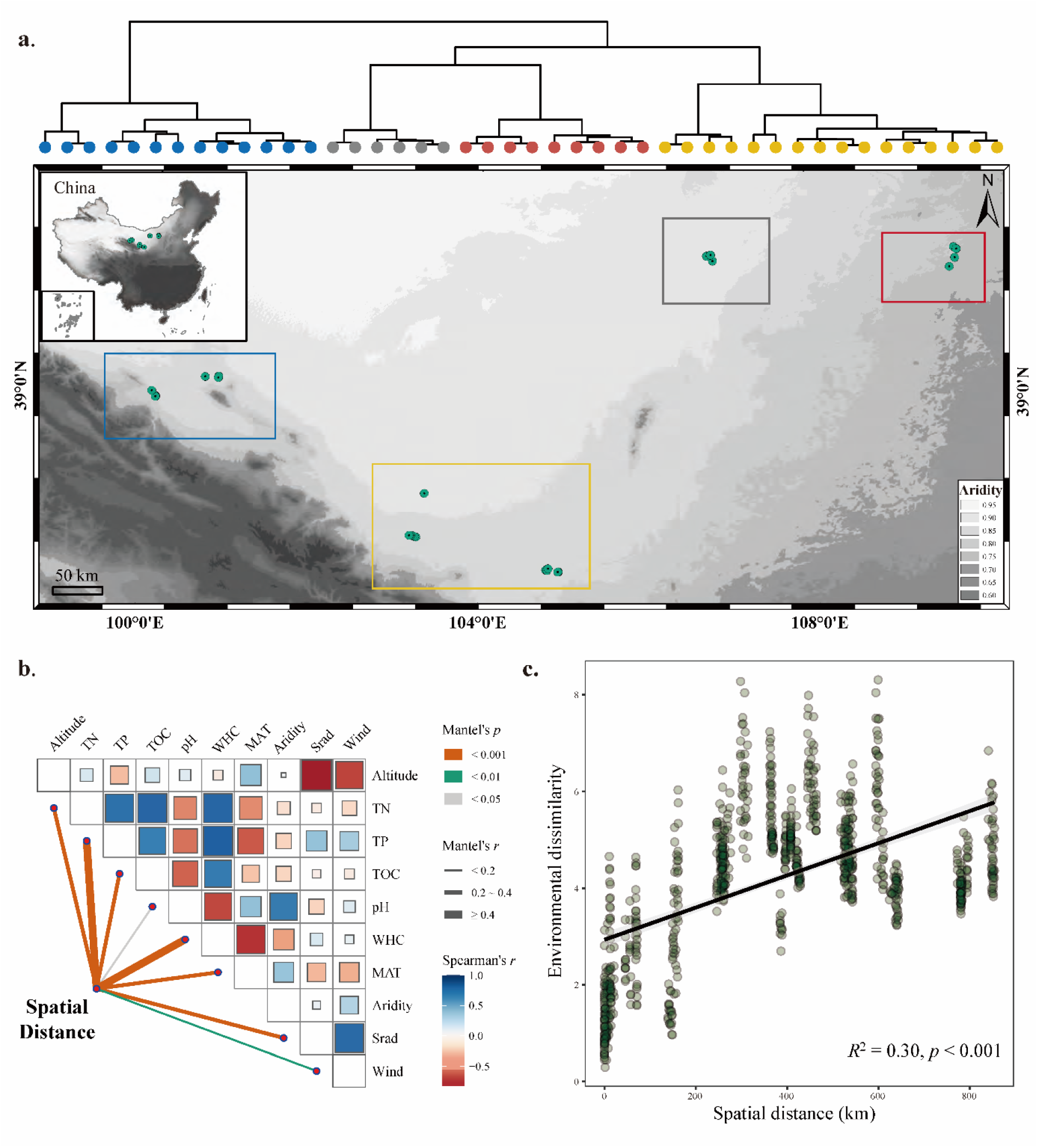
The location of sampling sites in Northern China and the relationships between environmental factors and spatial distance. The map of sampling sites, in which green points represent sampling sites with a gray gradient of Aridity (1-aridity index); and the clustering tree of sites is plotted based on the dissimilarity of environmental factors (**a**). Mantel test exhibits Spearman’s correlation coefficients between each pair of spatial distance and abiotic variables (**b**). Linear regression analysis evaluates the linkage of spatial distance with environmental dissimilarity (**c**).

### 2.2 Estimating environmental factors

A set of 10 key abiotic variables was estimated, including altitude, five local edaphic variables, and four regional climatic variables (**Table S2**). Spatial data (elevation, longitude, and latitude) was recorded by a handled GPS device in the field. Local edaphic variables included total nitrogen (TN), total phosphorus (TP), total organic carbon (TOC), water-holding capacity (WHC), and soil pH. These variables were measured as described in **Supplementary Note 1**. The climatic variables, such as mean annual temperature (MAT, °C), solar radiation (Srad, kJ m^-2^ day^-1^), wind speed (Wind, m s^-1^), and aridity index (AI), were obtained from Worldclim and Global Aridity and Potential Evapotranspiration databases [42, 43]. To avoid the problem of multicollinearity (**Figure S1**), an index of 1-AI (Aridity) was retained in all analyses, which considered the integrated effect of both precipitation and evaporation. We evaluated the correlations between the dissimilarity of environmental factors and pairwise spatial distance of sites. Environmental dissimilarity was calculated as the Euclidean distance using a matrix of normalized variables (*Z*-scores). Spatial distance was determined as the straight-line geographic distance by using the *R* package geosphere [44]. In our dataset, each study region posed a particular property of habitat. The differences of either individual environmental variables or the synthetic index exhibited positive associations with the spatial distance between sites at the large scale (*p* < 0.05, **Figure 1b, c**).

### 2.3 Illumina sequencing and morphological analysis

For each sample, total genomic DNA was extracted from 10 g of defrosted soil using the DNeasy PowerMax Soil Kit (Qiagen, USA). We used a group-special primer set CYA359f (5′-GGG GAA TYT TCC GCA ATG GG-3′) and CYA781r (A: 5′-GAC TAC TGG GGT ATC TAA TCC CAT T-3′; B: 5′-GAC TAC AGG GGT ATC TAA TCC CTT T-3′) to amplify the V3-V4 regions of cyanobacterial 16S rRNA genes [45]. The high-throughput sequencing method was used to assess the composition of cyanobacterial communities on an Illumina MiSeq PE300 platform [46]. Sequences with a similarity level of 100% were clustered into a separate operational taxonomic unit (zero-radius OTU, hereafter species) [47], and the taxonomic information was annotated by RDP Classifier based on SILVA v132 database. Unclassified, non-cyanobacterial species and eukaryotic chloroplasts were declined from our dataset. We then randomly resampled the subset of sequences with 25019 reads from each sample to control the sequencing effort. The filtered dataset contained 1100836 reads assigned into 1167 cyanobacterial species (**Figure S2**). To gain conservative classifying of morphotypes, we just selected the species with definite taxonomic annotation at the genus level. Therefore, a table comprised of 613 cyanobacterial species was ultimately used to identify the morphological characteristics, which were clustered into four aforementioned types, based on the detailed description of genera in AlgaeBase and CyanoDB 2 databases [48, 49].

### 2.4 Community trait analyses

The taxonomy-assigned species were used to construct the maximum likelihood phylogenetic trees for each morphotype with 1000 bootstrapped replicates in IQ-TREE v1.6.2 (http://www.iqtree.org/). The best model for building individual trees was evaluated by comparing the models’ Bayesian Information Criterion scores (BICs). Local *α*-diversity was measured as the observed number of species (richness). We summarized the abiotic features of sites by extracting the top three rotated principal component scores (RCs) of all environmental variables, conducted by the *R* package psych (**Table S3**). The correlations between the RCs and the relative abundance of each species were estimated based on Spearman’s coefficients, and niche differences thereby were evaluated. Then, we tested the phylogenetic signal by relating niche difference and phylogenetic distance in the *R* package vegan. In brief, we calculated abundance-weighted mean nearest taxon distance (*β*MNTD) and Bray-Curtis dissimilarity as the metrics of between-community phylogenetic and taxonomic *β*-diversity, respectively. The relative contributions of deterministic (heterogeneous or homogenous selection) and stochastic processes (dispersal limitation, homogenizing dispersal and ecological drift) for community assembly were calculated based on the null model to generate *β*-nearest taxon index (*β*NTI) and Raup-Crick index (RC_bray_) [14]. The neutral model was also constructed to determine the potential importance of stochastic processes and eatimate the migration rates (*m-*value) [50]. The detailed methods of modeling ananlyses are available in **Supplementary Note 2**.

We conducted random forest analysis in the *R* packages randomForest and rfPermute to evaluate the importance of each environmental factor on *β*NTI. To address multicollinearity problems, Spearman’s rank correlation coefficient (*ρ*) was measured between all pairs of variables, and high correlated factors (|*ρ*| > 0.60) were prevented from being invoked in one analysis. We used the linear regression to evaluate the relationships between *β*-diversity (Bray-Curtis dissimilarity, *β*MNTD) and normalized spatial distance, as well as between assembly processes (*β*NTI) and the variation of edaphic and climatic factors for each cyanobacterial subcommunity. The significant differences between regression slopes (*p*-value) were checked in the *R* package simba. We performed Wilcoxon or Kruskal-Wallis test to compare the differences of *α*-/*β*-diversity and relative abundance among four morphotypes. The contributions of environmental dissimilarity and spatial distance on *β*-diversity were quantified by regular and partial Mantel tests.

## 3. Results

### 3.1 Composition of cyanobacterial morphotypes

In the entire metacommunity, a total of 15 cyanobacterial genera were identified comprised of 613 relatively taxonomy-definite species across the regions, which were categorized into four morphotypes (**Table 1, Table S4**). We found that the different morphotypes were characterized by significant variations of relative abundance and species richness (**Figure 2a, Table S5**). The bundle-forming group was the most dominant morphotype (41.04% of sequencing reads in 277 species), followed by other non-heterocyst filamentous (16.24% in 147 species), heterocystous (9.70% in 119 species), and unicellular/colonial groups (4.36% in 70 species). It had a higher level of *α*-diversity (richness) than other morphotypes (Kruskal-Wallis test, *p* < 0.001). As expected, by estimating the occurrence frequency of each species, bundle-forming cyanobacteria occurred more widely in drylands, while a considerable proportion of non-heterocyst filamentous, heterocystous, and unicellular/colonial types preferred relatively special habitats limited to a few sites (**Figure 2b**). The top 15 abundant species were morphologically composed of nine bundle-forming, three non-heterocyst filamentous, one heterocystous, and two unicellular/colonial species, wherein *Microcoleus* sp. was the most abundant genus representing an average of 19.46% of sequencing reads in all sites (**Figure 2d**).

**Table 1.**
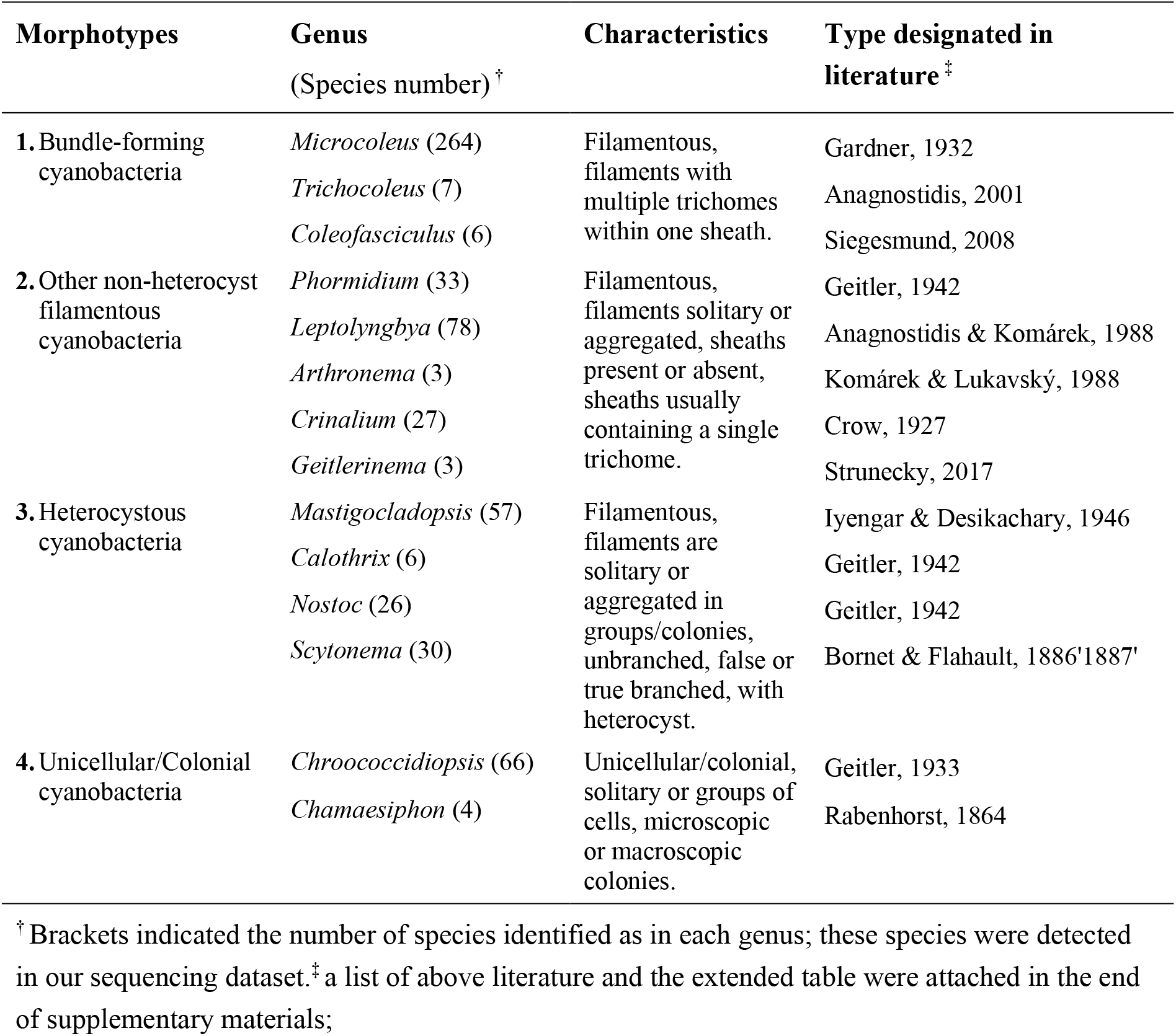
Summary of identified terrestrial cyanobacterial genera assigned into four main morphotypes

**Fig 2.**
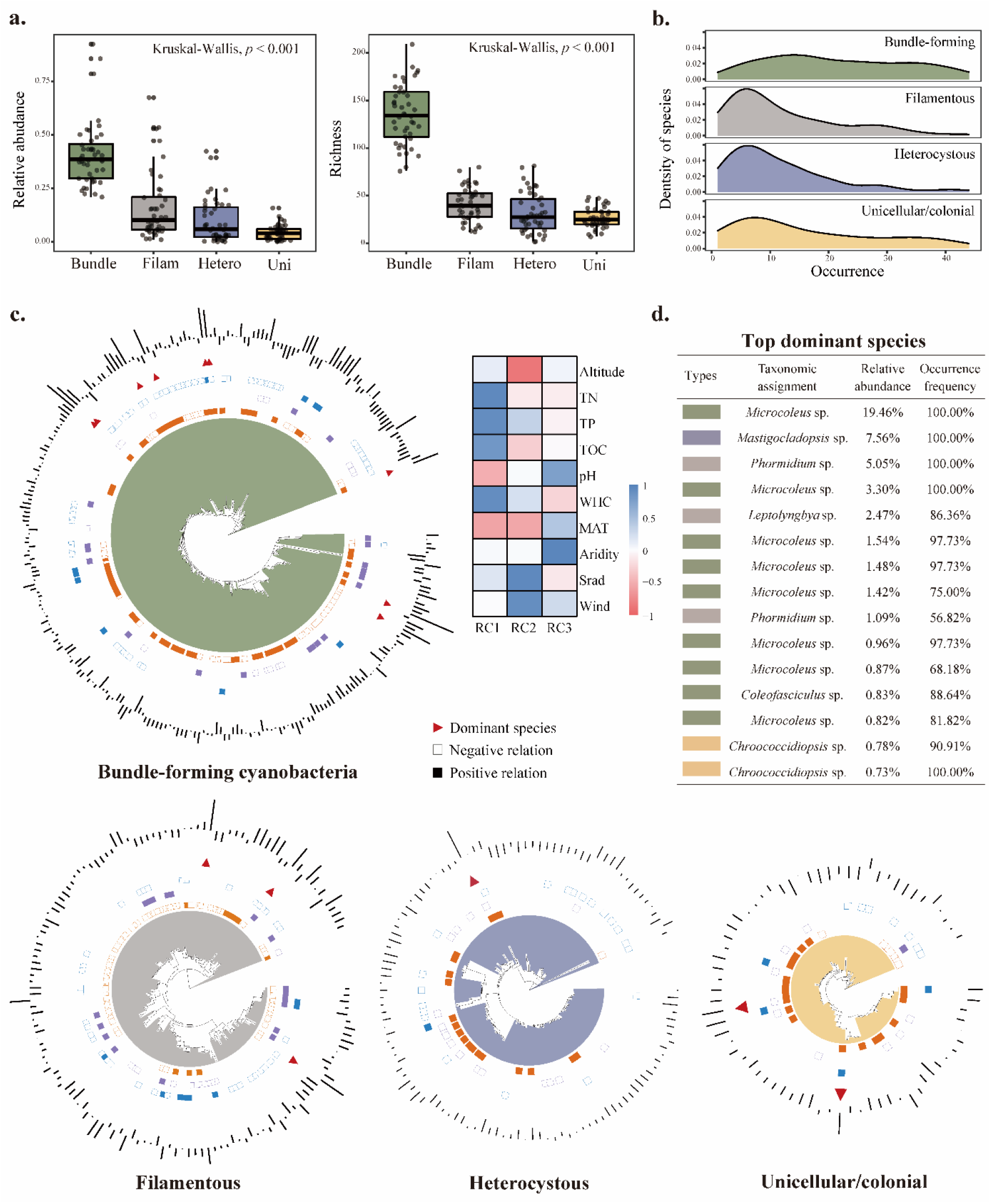
Community traits and phylogenetic distribution with ecological preferences of cyanobacterial species. Relative abundance and richness of different morphotypes (**a**), as well as species density along with the increase of occurrence frequency (**b**), are shown. The circle heatmaps outside phylogenetic trees of four morphotypes represent the positive or negative correlations between three rotated principal components (RCs) of environmental variables with the relative abundance of each taxon. The loadings of each environmental variable on RCs are exhibited behind. The bar chart around each phylogenetic tree shows the relative abundance (log-transformed). The outer bars are greater than the mean value of abundance, and the inner bars are less than it (**c**). The positions of the top 15 abundant taxa are marked as red triangles, and their taxonomic assignments are shown in the right table (**d**).

We then examined the linkages of cyanobacterial relative abundance and richness to environmental factors. The results suggested that TN and TOC contents were the most important factors generally related to species abundances of non-heterocyst filamentous, heterocystous, and unicellular/colonial cyanobacteria (*p* < 0.05), whilst the bundle-forming type was independent of all measured factors (**Figure S3**). Although the ecological preference of various species on environmental conditions was discrepant more or less, *Microcoleus* spp., as the widespread dominant bundle-forming cyanobacteria, consistently showed poor links with environmental dissimilarity (*i*.*e*., RCs, **Figure 2c**). Nonetheless, soil pH, WHC, and MAT were significantly related to the richness of bundle-forming type. In contrast, a set of abiotic factors, including TN, TP, TOC, WHC, and MAT, jointly impacted the abundance and richness of non-heterocyst filamentous cyanobacteria in biocrusts, indicating their sensitivity to environmental fluctuation.

### 3.2 Cyanobacterial *β*-diversity

Four morphological subcommunities of terrestrial cyanobacteria exhibited significant differences in *β*-diversity (Kruskal-Wallis test, *p* < 0.001, **Figure 3a**). We found that non-heterocyst filamentous cyanobacteria achieved the highest taxonomic (Bray-Curtis dissimilarity) and phylogenetic diversity (*β*MNTD), while taxonomic diversity of heterocystous morphotype and phylogenetic diversity of bundle-forming morphotype were lower than others, respectively. (Wilcoxon test, *p* < 0.001, **Table S5**). A distance-decay relationship between *β*-diversity and spatial distance was observed in all four morphotypes that either Bray-Curtis dissimilarity or *β*MNTD increased along with larger geographic interval of sites (*p* < 0.001, **Figure 3b, c**). However, the comparison of linear slopes demonstrated that the rates of both taxonomic and phylogenetic turnovers in the bundle-forming group were lower than others (**Figure S4**). The results of Mantel and partial Mantel tests indicated that spatial distance was a stronger predictor of *β*-diversity for heterocystous and unicellular/colonial morphotypes, while environmental dissimilarity explained more variance for the bundle-forming type (**Table S6**).

**Fig 3.**
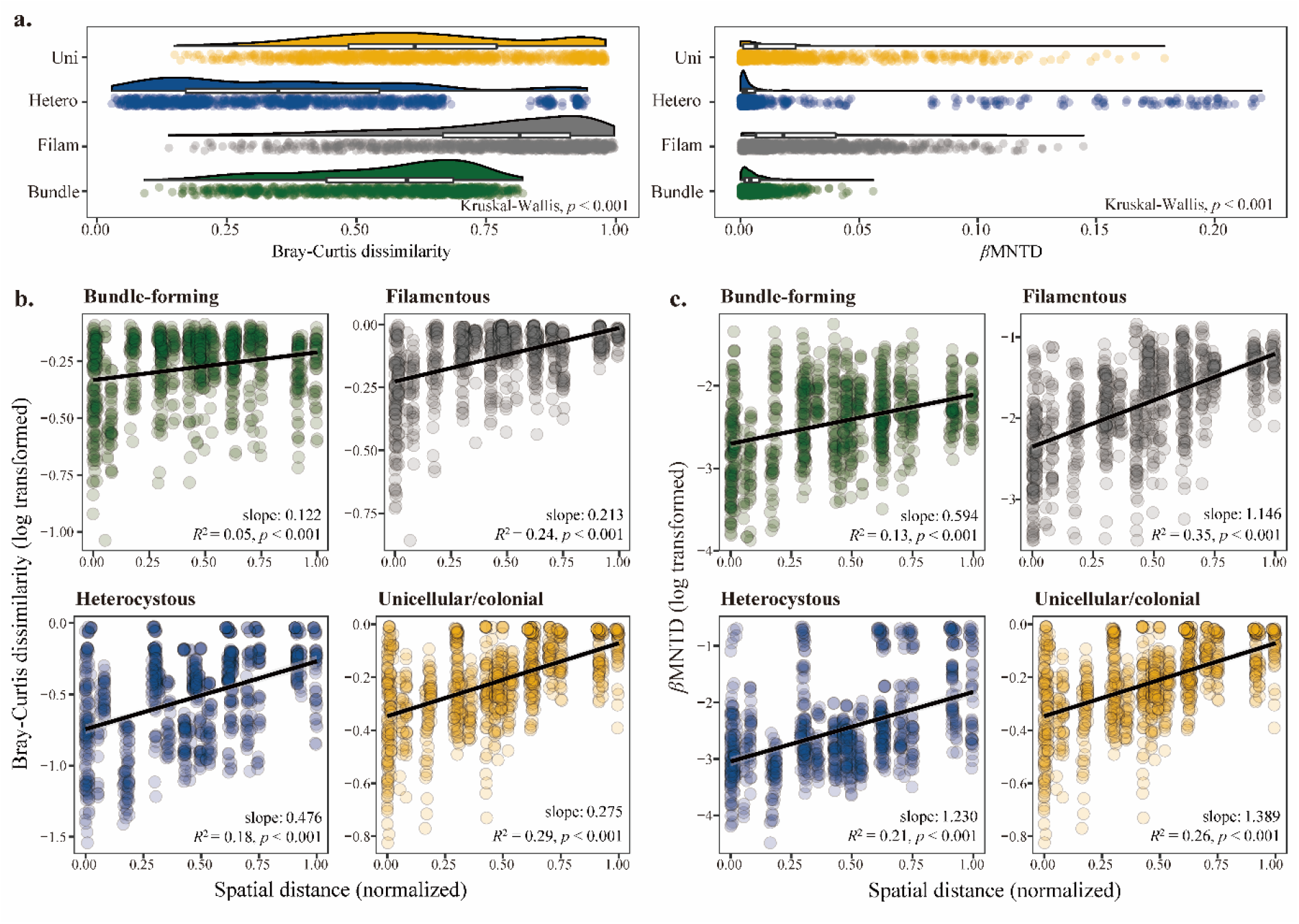
Geographical distribution of taxonomic and phylogenetic *β*-diversity of cyanobacterial morphotypes. Taxonomic and phylogenetic *β*-diversity are measured by Bray-Curtis dissimilarity and abundance-weighted *β*MNTD, respectively (**a**). The changes of taxonomic and phylogenetic *β*-diversity with the increase of spatial distance show positive relations in different morphotypes (**b, c**). Black lines represent significant linear regression fittings, and regression slopes, *R*^2^ values, and significant levels (*p*-value) are illustrated in the plots. The spatial distances are min-max normalized.

### 3.3 Community assembly processes

To select the most appropriate metrics for the assembly, we firstly compared the correlation coefficients between differences of environmental preference (RCs) and phylogenetic distance. A lot of phylogenetic signals were detected within relatively short phylogenetic distances in four morphotypes (adjusted *p* <0.05, **Figure S5**). Therefore, between-community MNTD (*i*.*e*., *β*MNTD) was measured to quantify the phylogenetic distance between each species in one community and its closest relative in a second community. We then used *β*NTI and RC_bray_ to check the relative influence of determinism and stochasticity on community assembly. The results revealed dominant roles of ecological drift (a mean value of 57.0% in all types, as a stochastic process) and heterogeneous selection (28.9%, as a deterministic process) in the assembly of terrestrial cyanobacteria (**Figure 4a**). In contrast, homogeneous selection showed little impact across morphotypes (0.2%). Interestingly, we found that homogenizing dispersal had a considerable influence just on the assembly of heterocystous cyanobacteria (8.9%), while the other morphotypes usually exhibited the importance of dispersal limitation (14.8%). Besides, Sloan’s neutral model explained 37.6%, 26.1%, 37.5%, and 52.1% of variances in four morphotypes, respectively (**Figure 4b**). It suggested that stochastic processes, as well as deterministic ones, equally drove the assembly. High immigration rates were determined in bundle-forming and heterocystous cyanobacteria (*m* = 0.055), rather than in unicellular/colonial (*m* = 0.021) and other non-heterocyst filamentous groups (*m* = 0.007). This result was consistent with the analysis of the phylogenetic null model, which indicated fewer influences of dispersal limitation on bundle-forming and heterocystous cyanobacteria.

**Fig 4.**
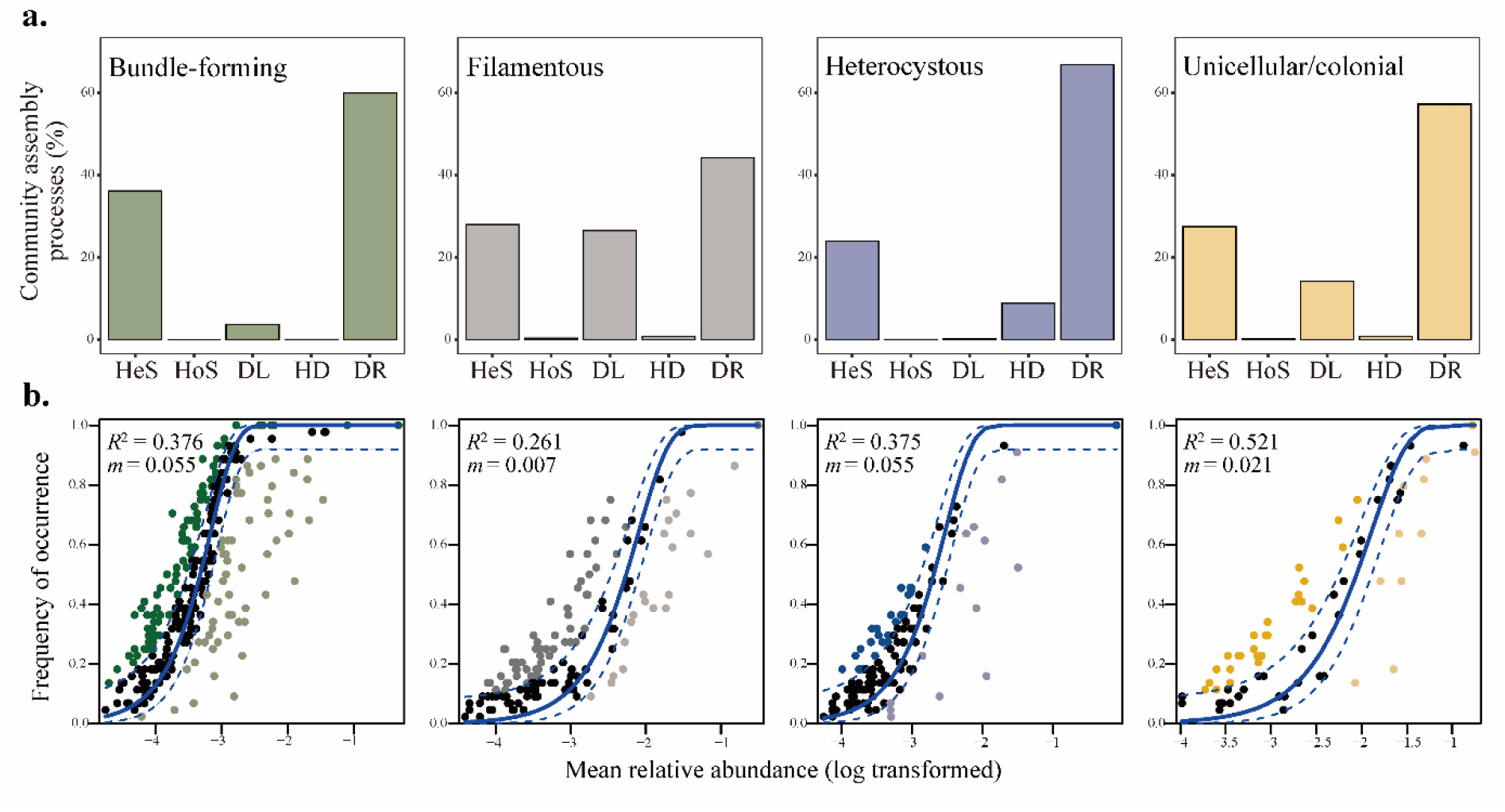
Assembly processes of different cyanobacterial subcommunities. The relative contributions of each process are estimated by the phylogenetic null model analysis (**a**). Five ecological processes, including heterogeneous selection (HeS), homogeneous selection (HoS), dispersal limitation (DL), homogenizing dispersal (HD), and ecological drift (DR), are invoked according to Vellend’s framework. Sloan’s abundance-based neutral model analysis on community assembly is further conducted (**b**). The *R*^2^ values indicate the overall fit of the neutral models, and the estimated *m*-value depicts the extent of immigration. The continuous blue lines represent the best-fitting neutral models, and the dashed ones represent the 95% confidence intervals.

We performed random forest analysis to find the key factors mediating the balance between determinism and stochasticity. The differences in spatial distance and WHC were excluded as explanatory variables due to multicollinearity problems (**Figure S6**). The value of *R*^2^ quantified the extent to which abiotic factors explain the total variance of *β*NTI. A normalized importance score of each predictor was assessed as the percentage of increase of mean squared error (MSE). In total, a range of 24.3∼52.7% variance in *β*NTI could be interpreted as the effect of abiotic factors on the community assembly. The results demonstrated that aridity difference (ΔAridity) was the most crucial variable for *β*NTI in all morphotypes (47.2% of mean increase in MSE, **Figure 5a**). Except for the heterogeneous group, MAT was an alternative important predictor, which had 31.9% of a mean increase of MSE, whereas TN just had a mean of 15.4% in different morphotypes. Then, we established linear regression between the change of abiotic factors and *β*NTI to further compare the difference of environmental responses among morphotypes (**Figure 5b**). The result is consistent with the ramdon forest analysis. Moreover, the pairwise significant increases of regression slopes informed a shifting threshold of ΔAridity influencing the assembly of different morphotypes, and the slope of bundle-forming cyanobacteria was particularly lower than the other groups. However, the other measured factors just had significant effects in one certain group. For example, the difference of nutritional conditions (ΔTN, ΔTP, and ΔTOC) controlled *β*NTI of bundle-forming and other non-heterocyst filamentous cyanobacteria, while climatic factors (ΔMAT, ΔSard, and ΔWind) were of vital importance on unicellular/colonial cyanobacteria.

**Fig 5.**
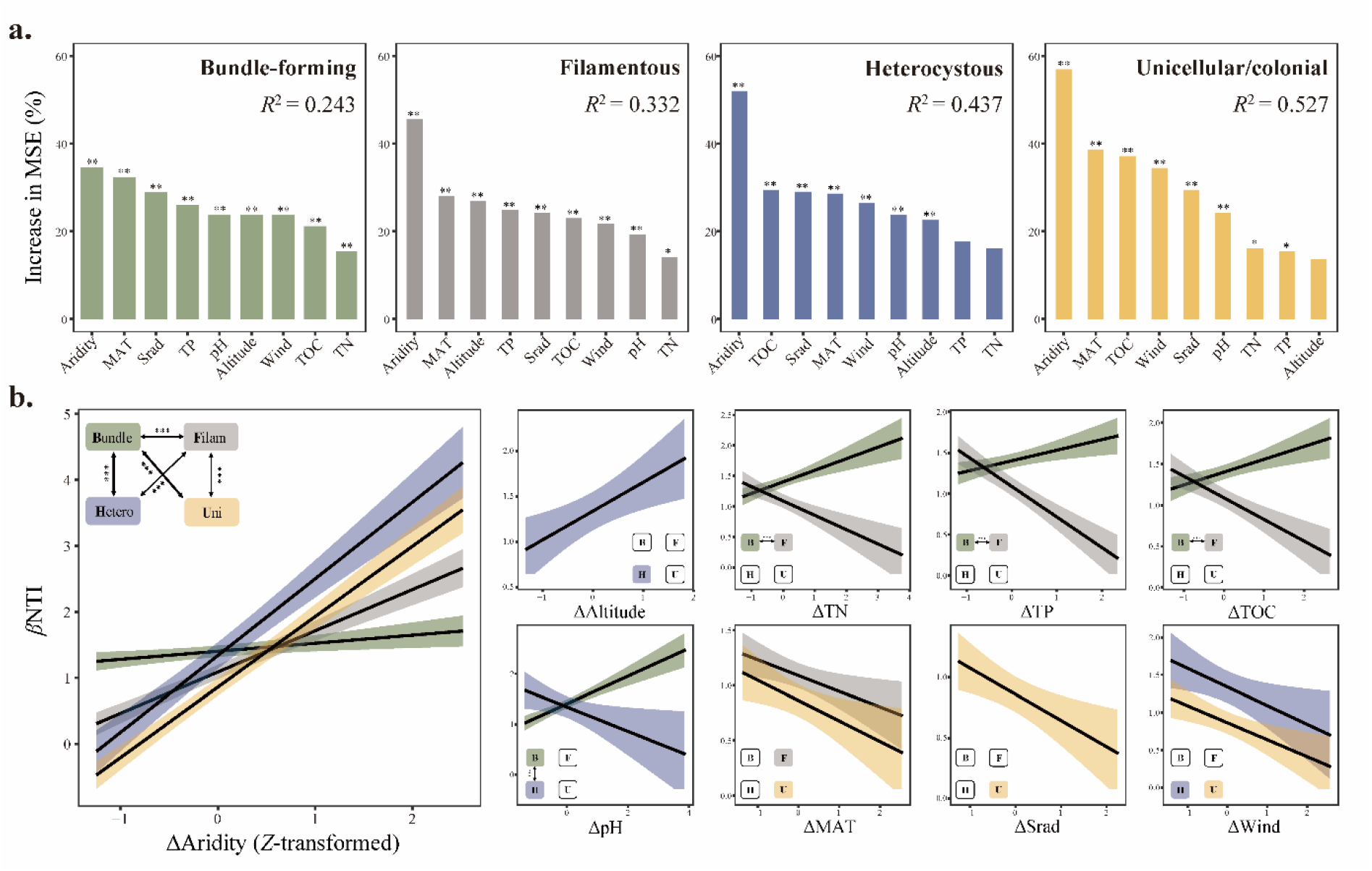
The effects of environmental variables on the assembly of cyanobacteria morphotypes. The relative importance of individual variables on *β*NTI is evaluated by random forest analysis. The value of *R*^2^ indicates the extent to which abiotic factors could explain the total variance of *β*NTI, and the asterisks represent the significant levels (**a**). Linear regressions between *β*NTI and the changes of environmental variables (*Z*-transformed) are built, and the black lines represent significant linear fittings with 95% confidence intervals (CIs). Different colors of CIs consist of four morphotypes (**b**). Only these slopes of statistically significant fittings are plotted, and double-headed arrows indicate the associations of slopes between different morphotypes. The thickness of the arrow is proportional to the extent of the slope difference. B: bundle-forming (Bundle); F: filamentous (Filam); H: heterocystous (Hetero); U: unicellular/colonial morphotype (Uni). ^*^ *p* < 0.05, ^**^ *p* < 0.01, and ^***^ *p* < 0.001.

## 4. Discussion

Despite the great potential of morphological traits in shaping microbial adaptation and functioning [51], the mechanisms by which microorganisms construct the biogeographic pattern and how various morphotypes assemble are still unclear. One of the challenging matters that impedes our understanding of these concerns is the limited observable characteristics among microbial taxa, especially when facing the mega-diverse microbiota in the nature [30]. In drylands, the community of terrestrial cyanobacteria provides a desirable model system to pursue the crucial but distinguishing roles of morphological subcommunities and their differential manners of assembly [37, 52].

Cyanobacterial morphotypes have a delicate vertical distribution in the layers of biocrusts and perform detailed functioning [33]. As the main member, bundle-forming cyanobacteria build rope-like bundles by grouping many trichomes in a common exopolysaccharide (EPS) sheath, which could bind soil particles to form the initial biocrust layer and improve the ability of water retention [39]. Similarly, other filamentous cyanobacteria further enhance soil stability and resistance against wind and rain erosion via filament winding [37]. In the deeper layer of biocrusts, specialized heterocystous cyanobacteria are capable of fixing atmospheric nitrogen to promote nutrient conditions [53]. For unicellular/colonial cyanobacteria, although we are lacking sufficient awareness of their ecological functions, some common genera (*e*.*g*., *Chroococcidiopsis*) usually proliferate in biocrusts from the extreme arid environments[54]. Previous research has demonstrated that the composition of the cyanobacterial community in biocrusts varies with the geographical gradient [55]. In our study regions, the *Microcoleus*-dominated bundle-forming morphotype is widely distributed but poorly linked with any measured environmental variables. In contrast, other filamentous, heterocystous, and unicellular/colonial cyanobacteria exhibit more sensitivity to the ambient conditions, especially local edaphic nutrients (*e*.*g*., TN and TOC). This to some extent explains why bundle-forming cyanobacteria usually act as the pioneer species and other morphotypes colonize subsequently in the harsh habitats of global drylands [25]. Meanwhile, it is noteworthy that environmental responses of the *Microcoleus* genus are often differential among species [55, 56]. Despite that their abundance exhibits independent with abitic factors, species richness and the number of shared species within the bundle-forming morphotype fluctuate along with the gradient of environmental variables. Therefore, we suppose a redundant *r*-strategy of these key cyanobacteria persisting in the harsh drylands, by which abundance of this particular functional group was maintained under within-group shift of different species.

Although there has been substantial evidence that morphological characteristics (*e*.*g*., leaf shape, body size, and root phenotype) could drive species distribution and *β*-diversity in biological communities [57, 58], this essential topic does not receive sufficient focus its merits in microbiota, especially given its critical functional roles in various ecosystems [2, 59]. Using cyanobacteria as the model community, we ascertained the heterogeneity of spatial turnover among microbial morphotypes within the phototrophic level. Despite that both taxonomic and phylogenetic *β*-diversity showed a distance-decay pattern in all types, the spatial turnover rate of bundle-forming cyanobacteria is particularly lower than others. Previous studies have suggested that the relative importance of environmental dissimilarity and spatial distance is largely related to the life history of a given microbial species (*e*.*g*., dispersal ability, arriving order, and fitness), and these factors jointly depict the pattern of spatial distribution [60, 61]. Thus, the low turnover rate of bundle-forming morphotype may result from strong adaptability and high dispersal. As proof, we indeed observed relative stability of abundance in the bundle-forming type against abiotic variation. The broader environmental breadth is derived from their prominent ability to excrete a mass of EPSs surrounding the trichomes and their capability of gliding locomotion within the EPS sheath, which gives them an ingenious behavior to escape the surge of environmental stresses (*e*.*g*., UV radiation, drought and wind erosion) [39, 62]. On the other hand, the phylogenetic null model and neutral model analyses indicated that stochastic processes, especially ecological drift, governed the assembly of the whole community, inferring possible relevancy between species dispersal and abundance. The global biogeographical pattern of bacteria has been verified that the cosmopolitan taxa are also the most abundant in individual assemblages because higher local population sizes could enable a wider potential of dispersal [63, 64]. The neutral model provides evidence that bundle-forming and heterocystous morphotypes possess the highest rate of immigration (*m*-value), rather than filamentous and unicellular/colonial types. Together, it implies that the strong adaptability and dominant abundance help bundle-forming cyanobacteria to achieve extensive distribution via air-borne diffusion at the cross-regional even global scale [65].

It is a still-controversial issue whether determinism or stochasticity contributes more to shaping microbial communities [17, 66]. Although our results suggested that the stochastic process is the primary factor in the assembly of cyanobacteria, heterogeneous selection, as a deterministic process, equally accounts for a considerable fraction of community variances. However, the balance of influences between deterministic and stochastic processes can be regulated by local edaphic (*e*.*g*., pH, salinity, nutrients) and climatic conditions [19, 23, 67]. For terrestrial cyanobacteria, we found that aridity is a nonnegligible driver on the changes of *β*NTI, as the indicator of community assembly. Precipitation has been supposed to strongly affect the global distribution of soil microorganisms by controlling soil moisture, which dictates the maintenance of ecosystem functions and the primary succession in drylands [3]. In the cross-regional scale, heterogeneous selection tends to be more dominant in the assembly of four cyanobacterial morphotypes along with the increase of ΔAridity, but this tendency is much weaker in bundle-forming than heterocystous or unicellular/colonial cyanobacteria. It demonstrates differential responses of various morphotypes in the assembly of subcommunities under climate change.

In conclusion, we inspected whether and to which extent morphological traits of terrestrial cyanobacteria can decide their biogeographic distributions and community assembly. The bundle-forming cyanobacteria are dominant at the cross-regional scale and show a lower rate of spatial turnover with remarkable independence against abiotic variables. The assembly is jointly governed by stochastic and deterministic processes, but aridity modulates a shifting weightiness among different morphotypes. The bundle-forming cyanobacteria are controlled by stochasticity with the change of aridity, while the assembly of heterocystous cyanobacteria turns from being relatively stochastic to deterministic at a lower threshold of ΔAridity. It has to be acknowledged that how important are morphological traits of microbes in maintaining soil biodiversity and ecosystem functioning is an ongoing attractive question. It deserves new biotechnologies to advance trait-based ecological studies in a changing mega-diverse world.

## Acknowledgments

The research is supported by the National Natural Science Foundation of China (Grant No. 41877419). The sequencing analysis is assisted by the Supercomputing Center of CAS, Wuhan Branch. We are grateful for technical support from the Analysis and Testing Center of IHB and for map plotting by Xingcheng Wang. The authors have no conflict of interest to declare.

## Data Accessibility Statement

The raw dataset is deposited in Dryad (https://doi.org/10.5061/dryad.95x69p8n3). MiSeq sequences are available in the Sequence Read Archive of the National Center for Biotechnology Information (NCBI-SRA, https://www.ncbi.nlm.nih.gov/sra/docs/) under the accession numbers SRR18056183 through SRR18056227. The climatic datasets are obtained from the WorldClim database (http://www.worldclim.org/).

