## Supplementary for "Aridity modulates biogeographic distribution and community assembly of cyanobacterial morphotypes in drylands"

**Author for contact**

* Dr. Hua Li (Institute of Hydrobiology, Chinese Academy of Sciences)

Address: South Donghu Road 7, 430072 Wuhan, China.

**Supplementary information** contains:

**Notes’** number: **2**

**Figures**’ number: **6**;

**Tables**’ number: **6**.

**Supplementary Note 1** Edaphic variables measurement.

**Supplementary Note 2** Analyses of community assembly.

**Supplementary Figure 1** The linkage between annual mean precipitation and aridity index based on linear regression.

**Supplementary Figure 2** Species rarefaction and accumulation curves at the local level and across the regional scale (n=45). The identification of cyanobacterial species is based on the similarity level of 100% threshold (One sample was excluded in the following steps owing to insufficient sequencing depth).

**Supplementary Figure 3** The relationships between abundance, richness, and environmental factors based on Spearman’s correlation. The color gradient represents the value of the correlation coefficient. The significant levels are denoted by asterisks, as * ($p < 0.05$), ** ($p < 0.01$) and *** ($p < 0.001$).

**Supplementary Figure 4** The differences of regression slope between spatial distance and $\beta$-diversity. The significant levels ($p$-value) of the linear regression are indicated as * ($p < 0.05$), ** ($p < 0.01$), *** ($p < 0.001$) and n.s. (non-significant).

**Supplementary Figure 5** Phylogenetic signal test based on relating species niche differences to phylogenetic distances using Mantel correlogram (Spearman's $r$-value). Solid and open symbols represent the significant and non-significant correlations.

**Supplementary Figure 6** Relationships between the differences of variables. The lower triangular matrix presents the pairwise linear relationships between soil variables; the upper triangular matrix presents Spearman's correlation coefficients of each fitting; the significant levels ($p$-value) of the linear regression are indicated as * ($p<0.05$), ** ($p<0.01$), and *** ($p<0.05$).

**Supplementary Table 1** The location of sampling sites in our study (n = 45).

**Supplementary Table 2** Summary of environmental variables in the study sites (n = 44).

**Supplementary Table 3** The standardized loadings of environmental factors on each principal component (n = 10).

**Supplementary Table 4** The detailed assignment of cyanobacterial morphotypes. The range and mean relative abundance, as well as species richness, of each subcommunity are shown in the brackets.

**Supplementary Table 5** Pairwise comparisons of cyanobacterial abundance, richness, and $\beta$-diversity (Bray-Curtis dissimilarity and $\beta$MNTD) between cyanobacterial morphotypes using the Wilcoxon test. The four morphotypes, bundle-forming, other non-heterocyst filamentous, heterocystous, and unicellular/colonial cyanobacteria are denoted by Bundle, Filam, Hetero, and Uni.

**Supplementary Table 6** Mantel and partial Mantel tests for the correlations between $\beta$-diversity (Bray-Curtis dissimilarity and $\beta$MNTD) and the explanatory variables (spatial and environmental distance, indicated by *Geodist* and *Envdist*) using Spearman's correlation coefficient.

**Supplementary Note 1 | Edaphic variables measurement.** The measurements of TN and TP were performed according to the standard methods [1]. To measure soil pH, subsamples were first suspended in centrifuge tubes with 1M KCl (1:5 ratio, w/w), then vortexed for 1 min and shaken for 30 min at room temperature. After a 60 min standing of the extract, the clear supernatant was filtered through a 0.45 μm Millipore filter and measured using a pH meter (Sartorius Intec, Germany) [2]. The combustion oxidation nondispersive infrared absorption method was used to determine TOC on a Vario TOC/$TN_b$ Select analyzer (Elementar, Germany). To measure WHC, 5 g of air-dried sample was thoroughly saturated with a known amount of water and then left to drain in a perforated centrifugal tube until the last drop of water had drained. Then, the value of WHC was calculated as the percent weight of water absorbed per gram of soil.

**Supplementary Note 2 | Analysis of community assembly.** Community assembly processes were estimated with a framework of phylogenetic null model, considering the criteria of taxonomic and phylogenetic turnovers within each pair of local cyanobacterial assemblages. Firstly, phylogenetic signal was tested by relating niche difference and phylogenetic distance. Niche differences of each species were calculated according to the orthogonal rotation of the top three scores (RCs), and between-species phylogenetic distances were measured in the *R* package iCAMP. We tested the phylogenetic signal of niche differences using the *R* package vegan to detect whether the ecological preference of a given cyanobacterial taxon is linked with phylogeny, and the level of significance (*p*-value) was adjusted by Bonferroni correction with 9999 permutations.

After that, we used the package iCAMP to calculate abundance-weighted mean nearest taxon distance ($\beta$MNTD) and Bray-Curtis dissimilarity as the metrics of between-community phylogenetic and taxonomic $\beta$-diversity, respectively [3, 4]. To quantify the degree to which observed values deviated from the distribution generated by a stochastic simulation, $\beta$-nearest taxon index ($\beta$NTI) and Raup-Crick index ($RC_{bray}$) were obtained by randomization (999 permutations) to null modeling on $\beta$MNTD (algorithm 'taxa shuffle') and Bray-Curtis dissimilarity ('frequency') [5]. A value of

|$\beta$NTI| > 2 was interpreted as deterministic processes that primarily govern observed turnover in pairwise comparisons of subcommunities, therein $\beta$NTI > 2 or < -2 further indicated that heterogeneous or homogenous selection was dominant significantly. In turn, a pair of subcommunities with |$\beta$NTI| < 2 meant that turnovers were governed by stochastic processes, such as dispersal limitation ($RC_{bray}$ > 0.95), homogenizing dispersal ($RC_{bray}$ < -0.95) and ecological drift (|$RC_{bray}$| < 0.95) [5]. We further constructed a neutral model to determine the potential importance of stochastic processes by predicting the relationship between species occurrence frequency and their relative abundance in the metacommunity [6]. The parameter $R^2$ presented the overall fit to the neutral model, and higher $R^2$ (> 0.2) indicated that the stochastic process of ecological drift and dispersal contributes more to the community assembly [7]. In addition, the inferred *m*-value was used to estimate migration rates, and higher *m* implied less dispersal limitation.

**Supplementary Figure 1 |** The linkage between annual mean precipitation and aridity index based on linear regression.

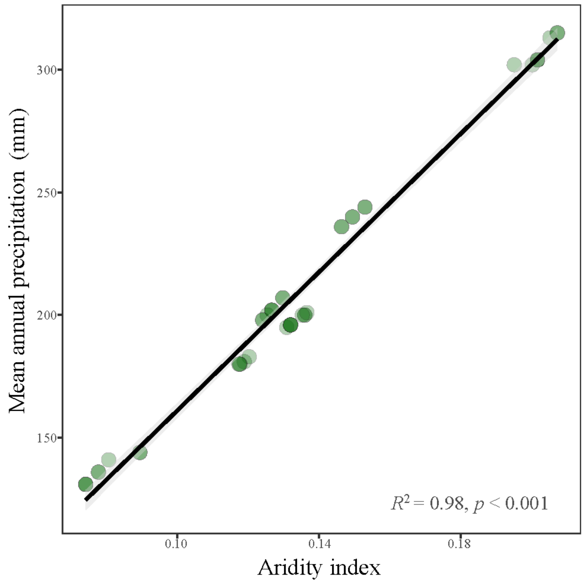

**Supplementary Figure 2 |** Species rarefaction and accumulation curves at the local level and across the regional scale (n=45). The identification of cyanobacterial species is based on the similarity level of 100% threshold (One sample was excluded in the following steps owing to insufficient sequencing depth).

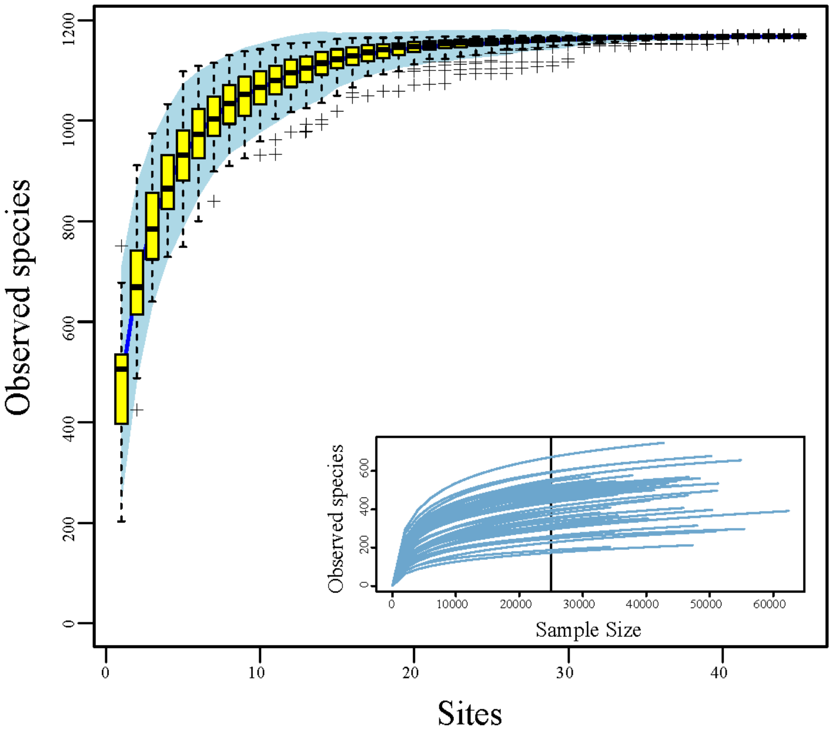

**Supplementary Figure 3 |** The relationships between abundance, richness and environmental factors based on Spearman's correlation. The color gradient represents the value of correlation coefficient. The significant levels are denoted by asterisks, as * ($p < 0.05$), ** ($p < 0.01$) and *** ($p < 0.001$).

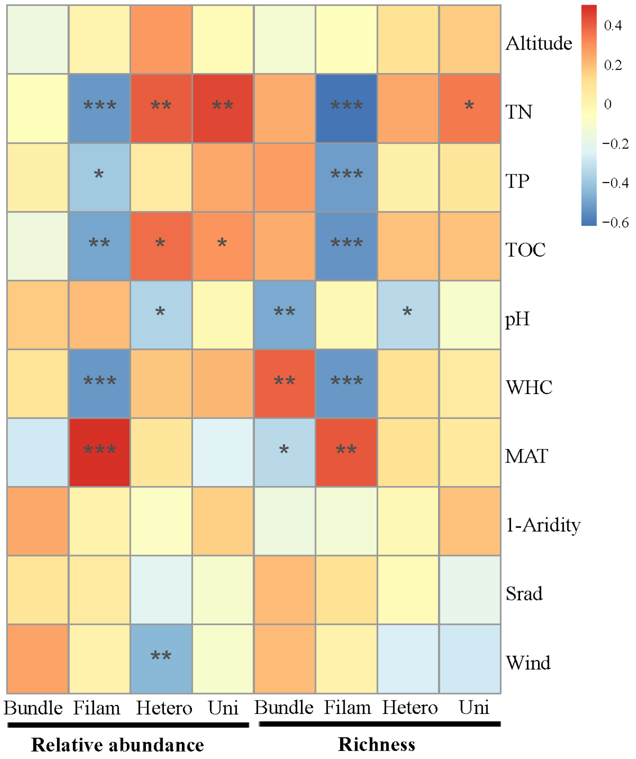

**Supplementary Figure 4 |** The differences of regression slope between spatial distance and *β*-diversity. The significant levels (*p*-value) of the linear regression are indicated as * ($p < 0.05$), ** ($p < 0.01$), *** ($p < 0.001$) and n.s. (non-significant).

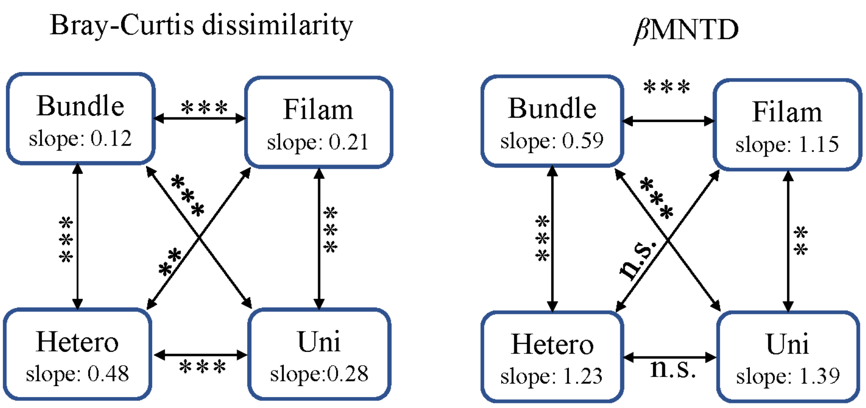

**Supplementary Figure 5** | Phylogenetic signal test based on relating species niche differences to phylogenetic distances using Mantel correlogram (Spearman's *r* value). Solid and open symbols represent the significant and non-significant correlations.

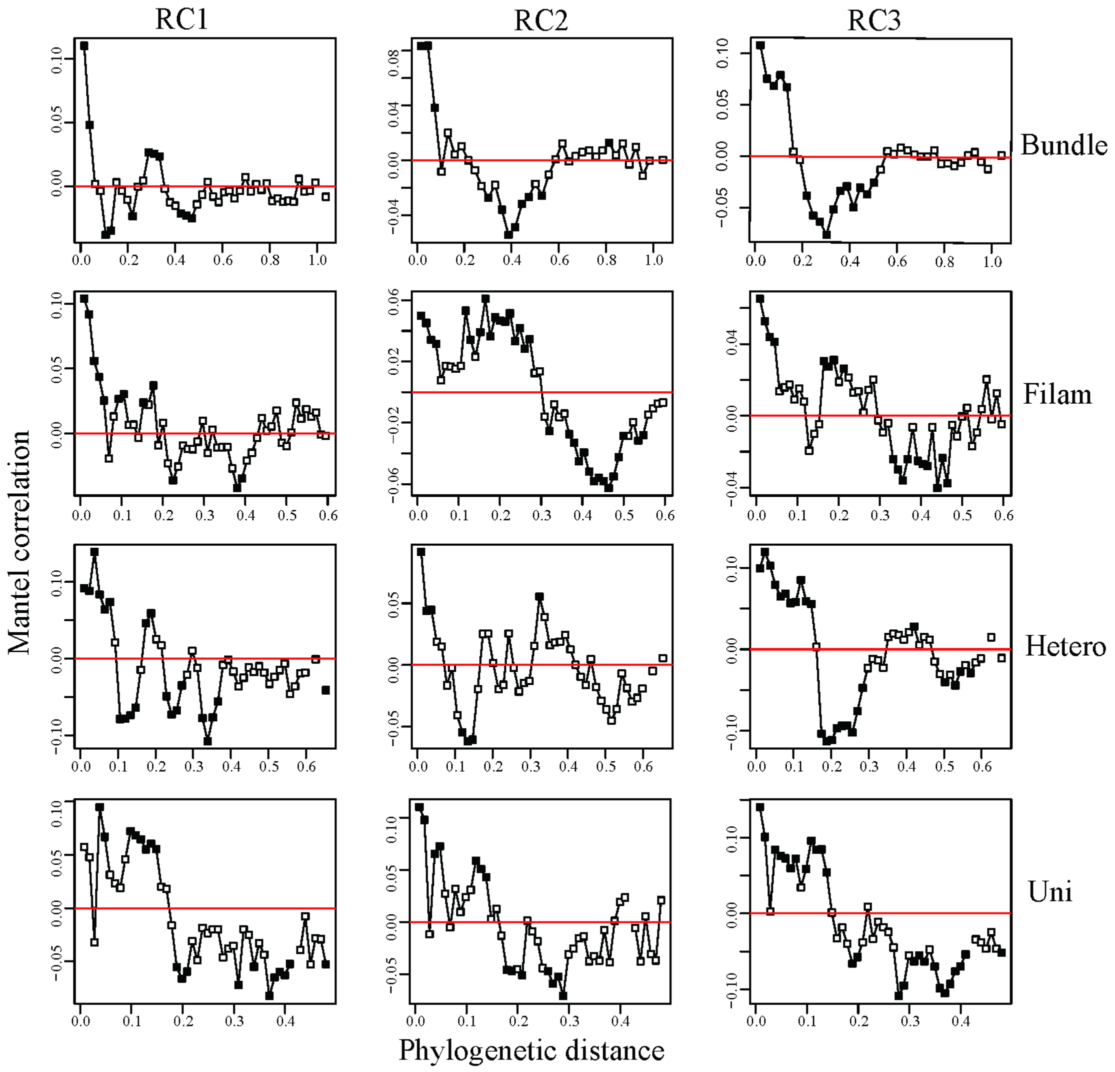

**Supplementary Figure 6 |** Relationships between the differences of variables. The lower triangular matrix presents the pairwise linear relationships between soil variables; the upper triangular matrix presents Spearman's correlation coefficients of each fitting; the significant levels (*p*-value) of the linear regression are indicated as * ($p$<0.05), ** ($p$<0.01), and *** ($p$<0.05).

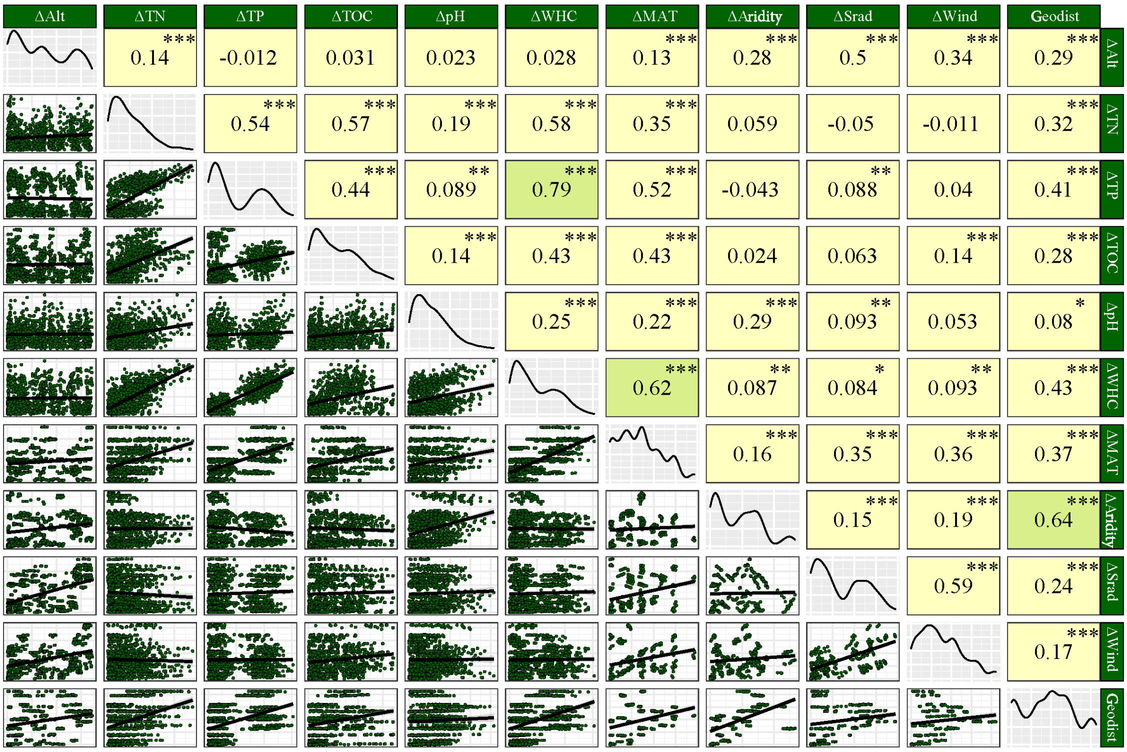

**Supplementary Table1 |** The location of sampling sites in our study (n = 45).

| | Sites | Longitude (°E) | Latitude (°N) | | Sites | Longitude (°E) | Latitude (°N) |
|---|---|---|---|---|---|---|---|
| ***Baituliang*** | BTL1 | 109.8126 | 40.3877 | ***Jiechaixian*** | JCX1 | 109.8181 | 40.2865 |
| | BTL2 | 109.8132 | 40.3876 | | JCX2 | 109.8179 | 40.2865 |
| | BTL3 | 109.8132 | 40.3872 | | JCX3 | 109.7471 | 40.2058 |
| | BTL4 | 109.8130 | 40.3878 | ***Nanhuzhen*** | NHZ1 | 103.3478 | 38.1686 |
| ***Dengkou*** | DK1 | 106.8382 | 40.3494 | | NHZ2 | 103.3478 | 38.1687 |
| | DK2 | 106.7645 | 40.3953 | ***Shahaowan*** | SHW1 | 104.7929 | 37.4638 |
| | DK3 | 106.7654 | 40.3947 | | SHW2 | 104.7945 | 37.4645 |
| | DK4 | 106.7668 | 40.3949 | | SHW3 | 104.8088 | 37.4596 |
| | DK5 | 106.8146 | 40.4045 | | SHW4 | 104.8218 | 37.4762 |
| | DK6 | 106.8152 | 40.4040 | ***Shapotou*** | SPT1 | 104.9421 | 37.4464 |
| ***Dalate*** | DLT1 | 109.8441 | 40.3667 | | SPT2 | 104.9410 | 37.4464 |
| | DLT2 | 109.8435 | 40.3667 | | SPT3 | 104.9404 | 37.4460 |
| ***Danxia*** | DX1 | 100.0852 | 38.9783 | | SPT4 | 104.9389 | 37.4468 |
| ***Geopark*** | DX2 | 100.0857 | 38.9777 | | SPT5 | 104.9360 | 37.4458 |
| | DX3 | 100.0856 | 38.9741 | ***Zhangye*** | ZY1 | 100.8323 | 39.1844 |
| | DX4 [§] | 100.0851 | 38.9729 | | ZY2 | 100.8320 | 39.1843 |
| ***Gaotai*** | GTX1 | 100.0379 | 39.0272 | | ZY3 | 100.8336 | 39.1849 |
| | GTX2 | 100.0379 | 39.0268 | | ZY4 | 100.8344 | 39.1977 |
| ***Huanghuatan*** | HHT1 | 103.2599 | 37.7596 | | ZY5 | 100.8338 | 39.1970 |
| | HHT2 | 103.2441 | 37.7633 | | ZY6 | 100.6715 | 39.1871 |
| | HHT3 | 103.2193 | 37.7705 | | ZY7 | 100.6719 | 39.1870 |
| | HHT4 | 103.1937 | 37.7725 | | ZY8 | 100.8319 | 39.1844 |
| | HHT5 | 103.1775 | 37.7732 | | | | |

[§] The site DX4 was exclude from subsequent analyses due to insufficient sequencing depth.

**Supplementary Table 2 |** Summary of environmental variables in the study sites (n = 44).

| Variables | Max | Min | Median | Mean | SD |
|---|---|---|---|---|---|
| **Altitude** (m) | 1876 | 1015 | 1557 | 1467 | 316 |
| **TN** (mg $g^{-1}$) | 2.02 | 0.10 | 0.65 | 0.72 | 0.43 |
| **TP** (mg $g^{-1}$) | 0.78 | 0.17 | 0.32 | 0.40 | 0.19 |
| **TOC** (mg $g^{-1}$) | 9.67 | 1.10 | 3.61 | 4.13 | 2.49 |
| **pH** (unitless) | 9.03 | 8.20 | 8.57 | 8.56 | 0.19 |
| **WHC** (%) | 52.32 | 21.81 | 31.01 | 33.28 | 8.20 |
| **MAT** (℃) | 9.24 | 5.37 | 7.33 | 7.38 | 1.17 |
| **Aridity** (unitless) | 0.93 | 0.79 | 0.87 | 0.86 | 0.04 |
| **Srad** (kJ $m^{-2}$ $day^{-1}$) | 16858 | 15691 | 16511 | 16362 | 373 |
| **Wind** (m $s^{-1}$) | 3.13 | 2.28 | 2.69 | 2.71 | 0.25 |
| **MAP** (mm) [§] | 315.00 | 131.00 | 200.00 | 212.66 | 56.12 |

[§] MAP and aridity Index (AI) are highly corelated. To avoid multicollinearity problem, only transformed Aridity (1-AI) is retained in the next analyses.

**Supplementary Table 3 |** The standardized loadings of environmental factors on each principal component (n = 10).

| **Variable** | **RC1** | **RC2** | **RC3** | **h2** | **u2** | **com** |
|---|---|---|---|---|---|---|
| Altitude | 0.12 | -0.86 | 0.07 | 0.75 | 0.246 | 1.1 |
| TN | 0.94 | -0.11 | -0.09 | 0.9 | 0.097 | 1 |
| TP | 0.91 | 0.34 | -0.06 | 0.95 | 0.05 | 1.3 |
| TOC | 0.83 | -0.29 | -0.02 | 0.78 | 0.224 | 1.2 |
| pH | -0.49 | 0.02 | 0.75 | 0.81 | 0.19 | 1.7 |
| WHC | 0.91 | 0.22 | -0.25 | 0.94 | 0.064 | 1.3 |
| MAT | -0.56 | -0.54 | 0.43 | 0.8 | 0.202 | 2.9 |
| Aridity | 0.03 | 0.02 | 0.97 | 0.95 | 0.054 | 1 |
| Srad | 0.17 | 0.93 | -0.12 | 0.9 | 0.101 | 1.1 |
| Wind | 0.01 | 0.9 | 0.27 | 0.89 | 0.115 | 1.2 |

**Supplementary Table 4 |** The detailed assignment of cyanobacterial morphotypes. The range and mean relative abundance, as well as species richness, of each subcommunities are shown in the brackets.

| **Morphotypes** | **Genus** |
|---|---|
| Bundle-forming cyanobacteria<br>(20.95~92.52%, mean 41.04%, 277 species) | *Microcoleus* [†] (11.51~92.5%, mean 39.16%)<br>*Trichocoleus* [†] (0~7.27%, mean 1.01%)<br>*Coleofasciculus* [†] (0~6.24%, mean 0.87%) |
| Other non-heterocyst filamentous cyanobacteria<br>(1.00~67.50%, mean 16.24%, 147 species) | *Phormidium* [†] (0.04~66.71%, mean 8.65%)<br>*Leptolyngbya* [†] (0~37.4%, mean 5.85%)<br>*Arthronema* [†] (0~10.61%, mean 0.65%)<br>*Crinalium* [†] (0~3.65%, mean 0.18%)<br>*Geitlerinema* [†] (0~5.11%, mean 0.53%)<br>*Planktothrix* [†, §] (0~9.26%, mean 0.39%) |
| Heterocystous cyanobacteria<br>(0.01~42.39%, mean 9.70%, 119 species) | *Mastigocladopsis* [†] (0~39.60%, mean 8.05%)<br>*Calothrix* [†] (0~0.48%, mean 0.01%)<br>*Nostoc* [†] (0~2.70, mean 0.77%)<br>*Scytonema* [†] (0-11.38%, mean 0.87%) |
| Unicellular/Colonial cyanobacteria<br>(0.07~15.80%, mean 4.36%, 70 species) | *Chroococcidiopsis* [†] (0.06~15.79%, mean 4.33%)<br>*Chamaesiphon* [†] (0~0.86%, mean 0.03%) |
| Unassigned fraction<br>(3.70~52.08%, mean 28.66%, 554 species) | Uncultured and unidentified to genus fractions |

[†] The detail morphological description can be found in the literatures, which can be found in the end of this supplementary; [§] *Planktothrix* is a common aquatic genus, thus an annotated error might be the case here.

**Supplementary Table 5 |** Pairwise comparisons of cyanobacterial abundance, richness and $\beta$-diversity (Bray-Curtis dissimilarity and $\beta$MNTD) between cyanobacterial morphotypes using the Wilcoxon test. The four morphotypes, bundle-forming, other non-heterocyst filamentous, heterocystous and unicellular/colonial cyanobacteria are denoted by Bundle, Filam, Hetero and Uni.

| | | **Abundance** | **Richness** | **Bray-Curtis** | **$\beta$MNTD** |
|---|---|---|---|---|---|
| Filam | Bundle | *** | *** | *** | *** |
| Hetero | Bundle | *** | *** | *** | *** |
| Uni | Bundle | *** | *** | *** | *** |
| Hetero | Filam | * | * | *** | *** |
| Uni | Filam | *** | *** | *** | *** |
| Uni | Hetero | * | n.s. | *** | *** |

The significant levels of the correlation are indicated as $^{***}$ ($p$<0.001), $^{**}$ ($p$<0.01), $^{*}$ ($p$<0.05) and n.s. (non-significant).

**Supplementary Table 6** Mantel and partial Mantel tests for the correlations between $\beta$-diversity (Bray-Curtis dissimilarity and $\beta$MNTD) and the explanatory variables (spatial and environmental distance, indicated by *Geodist* and *Envdist*) using Spearman's correlation coefficient.

| **Effects** | **Control for** | **Bundle** | | **Filam** | | **Hetero** | | **Uni** | |
|---|---|---|---|---|---|---|---|---|---|
| | | BC | $\beta$MNTD | BC | $\beta$MNTD | BC | $\beta$MNTD | BC | $\beta$MNTD |
| *Envdist* | | 0.198$^{***}$ | 0.448$^{***}$ | 0.454$^{***}$ | 0.305$^{***}$ | 0.092 | 0.133$^{*}$ | 0.193$^{**}$ | 0.157$^{**}$ |
| *Geodist* | | 0.173$^{**}$ | 0.338$^{***}$ | 0.427$^{***}$ | 0.605$^{***}$ | 0.397$^{***}$ | 0.451$^{***}$ | 0.534$^{***}$ | 0.493$^{***}$ |
| *Envdist* | *Geodist* | 0.137$^{*}$ | 0.353$^{***}$ | 0.325$^{***}$ | n.s. | n.s. | n.s. | n.s. | n.s. |
| *Geodist* | *Envdist* | 0.096$^{*}$ | 0.172$^{**}$ | 0.281$^{***}$ | 0.550$^{***}$ | 0.400$^{***}$ | 0.442$^{***}$ | 0.510$^{***}$ | 0.479$^{***}$ |

The significances are tested using 9999 permutations. Significant levels are indicated as $^{***}$ ($p$<0.001), $^{**}$ ($p$<0.01), $^{*}$ ($p$<0.05) and n.s. (non-significant).

**Extended Table.** Summary of typical terrestrial cyanobacterial genera assigned into the main morphotypes

| **Morphotypes** | **Genus** | **Characteristics** | **Type designated in literature** |
|---|---|---|---|
| **1.** Bundle-forming cyanobacteria<br>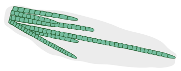 | *Microcoleus*<br>*Trichocoleus*<br>*Coleofasciculus*<br>*Schizothrix* | Filamentous, filaments with multiple trichomes within one sheath. | Gardner, 1932<br>Anagnostidis, 2001<br>Siegesmund, 2008<br>Gomont, 1892 |
| **2.** Other non-heterocyst filamentous cyanobacteria<br>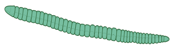 | *Phormidium*<br>*Leptolyngbya*<br>*Arthronema*<br>*Crinalium*<br>*Geitlerinema*<br>*Wilmottia*<br>*Oculatella*<br>*Phormidesmis*<br>*Oscillatoria*<br>*Porphyrosiphon*<br>*Pycnacronema*<br>*Potamolinea*<br>*Pseudophormidium*<br>*Komvophoron*<br>*Pantanalinema* | Filamentous, filaments solitary or aggregated, sheaths present or absent, sheaths usually containing a single trichome. | Geitler, 1942<br>Anagnostidis & Komárek, 1988<br>Komárek & Lukavský (1988)<br>Crow, 1927<br>Strunecky, 2017<br>Strunecký, 2011<br>Zammit, 2012<br>Turicchia, 2009<br>Gardner, 1932<br>Gomont, 1892<br>Martins, 2018<br>Martins & Branco, 2016<br>Anagnostidis & Komárek, 1988<br>Anagnostidis & Komárek, 1988<br>Vieira Vaz, 2015 |
| **3.** Heterocystous cyanobacteria<br>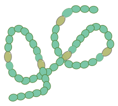 | *Mastigocladopsis*<br>*Calothrix*<br>*Nostoc*<br>*Scytonema*<br>*Tolypothrix*<br>*Mojavia*<br>*Macrochaete*<br>*Spirirestis*<br>*Brasilonema*<br>*Aetokthonos*<br>*Trichormus*<br>*Desmonostoc*<br>*Stigonema* | Filamentous, filaments are solitary or aggregated in groups/colonies, unbranched, false or true branched, with heterocyst. | Iyengar & Desikachary, 1946<br>Geitler, 1942<br>Geitler, 1942<br>Bornet & Flahault, 1886'1887'<br>Geitler, 1942<br>Reháková, 2007<br>Berrendero-Gómez, 2016<br>Flechtner, 2002<br>Fiore, 2007<br>Wilde, 2014<br>Komárek & Anagnostidis, 1989<br>Hrouzek, 2013<br>Gardner, 1932 |
| **4.** Unicellular/Colonial cyanobacteria<br>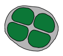 | *Chroococcidiopsis*<br>*Chamaesiphon*<br>*Pleurocapsa*<br>*Aliterella*<br>*Chlorogloea*<br>*Prochlorococcus*<br>*Chroococcus*<br>*Gloeocapsopsis*<br>*Aphanothece* | Unicellular/colonial, solitary or groups of cells, microscopic or macroscopic colonies. | Geitler, 1933<br>Rabenhorst, 1864<br>Hauck, 1885<br>Rigonato, 2016<br>Wille, 1900<br>Komárek, 2020<br>Nägeli, 1849<br>Komárek, 1993<br>Nägeli, 1849 |

166

### A list of literature in Table 1 (Extended)

Anagnostidis, K. (2001). Nomenclatural changes in cyanoprokaryotic order Oscillatoriales. Preslia, Praha 73: 359-375.

Anagnostidis, K. & Komárek, J. (1988). Modern approach to the classification system of cyanophytes. 3. Oscillatoriales. Archiv für Hydrobiologie, Supplement 80(1-4): 327-472.

Berrendero Gómez, E.B., Johansen, J.R., Kaštovský, J., Bohunická, M. & Capková, K. (2016). Macrochaete gen. nov. (Nostocales, Cyanobacteria), a taxon morphologically and molecularly distinct from Calothrix. Journal of Phycology 52(4): 638-655.

Bornet, É. & Flahault, C. (1886 '1887'). Revision des Nostocacées hétérocystées contenues dans les principaux herbiers de France (Troisième fragment). Annales des Sciences Naturelles, Botanique, Septième série 5: 51-129.

Crow, W.B. (1927). Crinalium, a new genus of Cyanophyceae, and its bearing on the morphology of the group. Annals of Botany 41: 161-166.

Fiore, M.F., Sant'Anna, C.L., de Paiva Azevedo, M. T., Komarek, J., Kaštovský, Sulek, J. & Lorenzi, A.S. (2007). The cyanobacterial genus Brasilonema, gen. nov., a molecular and phenotypic evaluation. Journal of Phycology 43: 789-798.

Flechtner, V.R., Boyer, S.L., Johansen, J.R. & DeNoble, M.L. (2002). Spirirestis rafaelensis gen. et sp. nov. (Cyanophyceae), a new cyanobacterial genus from arid soils. Nova Hedwigia 74: 1-24.

Gardner, N.L. (1932). The Myxophyceae of Porto Rico and the Virgin Islands. In: Scientific Survey of Porto Rico and the Virgin Islands. Volume VIII. Part 2. (Anon. Eds), pp. 249-311. New York: New York Academy of Sciences.

Geitler, L. (1933). Diagnosen neuer Blaualgen von den Sunda-Inseln. Arch. Hydrobiol. Suppl. 12: 622-634.

Geitler, L. (1942). Schizophyta: Klasse Schizophyceae. In: Die natürlichen Pflanzenfamilien, Sweite Auflage. (Engler, A. & Prantl, K. Eds) Vol.1b, pp. 1-232. Leipzig: Wilhelm Engelmann.

Gomont, M. (1892). Monographie des Oscillariées (Nostocacées homocystées). Annales des Sciences Naturelles, Botanique, Série 7 15: 263-368, pls 6-14 (VI-XIV in text and expl. pl.).

Hrouzek, P., Lukešová, A., Mareš, J. & Ventura, S. (2013). Description of the cyanobacterial genus Desmonostoc gen. nov. including D. muscorum comb. nov. as a distinct, phylogenetically coherent taxon related to the genus Nostoc. Fottea 13(2): 201-213.

Hauck, F. (1885). Die Meeresalgen Deutschlands und Österreichs. In: Kryptogamen-Flora von Deutschland, Österreich und der Schweiz. Zweite Auflage. (Rabenhorst, L. Eds) Vol. 2, pp. 513-575, [i]-xxiii [xxiv]. Leipzig: Eduard Kummer.

Iyengar, M.O.P. & Desikachary, T.V. (1946). Mastigocladopsis jogensis gen. et sp. nov., a new member of the Stigonemataceae. Proceedings of the Indian Academy of Sciences, Section B 24: 55-59.

Komárek, J. & Anagnostidis, K. (1989). Modern approach to the classification system of Cyanophytes 4 - Nostocales. Algological Studies 56: 247-345.

Komárek, J., Johansen, J.R., Smarda, J. & Strunecky, O. (2020). Phylogeny and taxonomy of Synechococcus-like cyanobacteria. Fottea 20(2): 171-191, 6 figures.

Komárek, J. (1993). Validation of the genera Gloeocapsopsis and Asterocapsa (Cyanoprokaryota) with regard to species from Japan, Mexico and Himalayas. Bulletin of the National Science Museum, Tokyo, Series B (Botany) 19(1): 19-37.

Komárek, J. & Lukavský, J. (1988). Arthronema, a new cyanophyte genus from Afro-Asian deserts. Algological Studies 50-53: 249-267.

Martins, M.D., Machado-de-Lima, N.M. & Branco, L.H.Z. (2018). Polyphasic approach using multilocus analyses supports the establishment of the new aerophytic cyanobacterial genus Pycnacronema (Coleofacisculaceae, Oscillatoriales). Journal of Phycology 55(1): 146-159.

Martins, M.D. & Branco, L.H.Z. (2016). Potamolinea gen. nov. (Oscillatoriales, Cyanobacteria): a phylogenetically and ecologically coherent cyanobacterial genus. International Journal of Systematic and Evolutionary Microbiology 66(9): 3632-3641, 3 fig., 2 tables.

Nägeli, C. (1849). Gattungen einzelliger Algen, physiologisch und systematisch bearbeitet. Neue Denkschriften der Allg. Schweizerischen Gesellschaft für die Gesammten Naturwissenschaften 10(7): i-viii, 1-139, pls I-VIII.

Rabenhorst, L. (1864). Flora europaea algarum aquae dulcis et submarinae. Sectio I. Algas diatomaceas complectens, cum figuris generum omnium xylographice impressis. pp. 1-359. Lipsiae [Leipzig]: Apud Eduardum Kummerum.

Reháková, K., Johansen, J.R., Casamatta, D.A., Xuesong, L., & Vincent, J. (2007). Morphological and molecular characterization of selected desert soil cyanobacteria: three species new to science including Mojavia pulchra gen. et sp. nov. Phycologia 46: 481-502.

Rigonato, J., Arantes Gama, W., Oliverira, D., Zanini Branco, L.H., Pereira Brandini, F., Genuário, D.B. & Fiore, M.F. (2016). Aliterella atlantica gen. nov., sp. nov., and Aliterella antarctica sp. nov., novel members of coccoid Cyanobacteria. International Journal of Systematic and Evolutionary Microbiology 66: 2853-2861, 4 figs.

Siegesmund, M.A., Johansen, J.R., Karsten, U. & Friedl, T. (2008). Coleofasciculus gen. nov. (Cyanobacteria): morphological and molecular criteria for revision of the genus Microcoleus Gomont. Journal of Phycology 44: 1572-1585.

Strunecký, O., Bohunická, M., Johansen, J.R., Čapková, K., Raabová, L., Dvořák, P. & Komárek, J. (2017): A revision of the genus Geitlerinema and a description of the genus Anagnostidinema gen. nov. (Oscillatoriophycidae, Cyanobacteria). Fottea, 17(1): 114–126

Strunecký, O., Elster, J. & Komárek, J. (2011). Taxonomic revision of the freshwater cyanobacterium "Phormidium" murrayi = Wilmottia murrayi. Fottea 11(1): 57-71.

Turicchia, S., Ventura, S., Komárková, J. & Komárek, J. (2009). Taxonomic evaluation of cyanobacterial microflora from alkaline marshes of northern Belize. 2. Diversity of oscillatorialean genera. Nova Hedwigia 89: 165-200, 26 figs., 1 table.

Vieira Vaz, M.G., Genuário, D.B., Andreote, A.P., Malone, C.F., Sant'Anna C.L, Barbiero, L. & Fiore, M.F. (2015). Pantanalinema gen. nov. and Alkalinema gen. nov.: novel pseudanabaenacean genera (Cyanobacteria) isolated from saline-alkaline lakes. International Journal of Systematic and Evolutionary Microbiology 65: 298-308, 4 figs.

Wille, N. (1900). Algologische Notizen I-VI. Nyt Magazin for Naturvidenskaberne 38(1): 1-27.

Wilde, S.B., J.R.Johansen, H.D.Wilde, P.Jiang, B.A.Bartelme & R.S.Haynie (2014). Aetokthonos hydrillicola gen. et sp. nov.: epiphytic cyanobacteria on invasive aquatic plants implicated in Avian Vacuolar Myelinopathy. Phytotaxa 181(5): 243-260, 7 figs, 4 tables.

Zammit, G., Billi, D. & Albertano, P. (2012). The subaerophytic cyanobacterium Oculatella subterranea (Oscillatoriales, Cyanophyceae) gen. et sp. nov.: a cytomorphological and molecular description. European Journal of Phycology 47(4): 341-354.
